# Stability and folding pathways of tetra-nucleosome from six-dimensional free energy surface

**DOI:** 10.1101/2021.01.06.425601

**Authors:** Xinqiang Ding, Xingcheng Lin, Bin Zhang

## Abstract

The three-dimensional organization of chromatin is expected to play critical roles in regulating genome functions. High-resolution characterization of its structure and dynamics could improve our understanding of gene regulation mechanisms but has remained challenging. Using a near-atomistic model that preserves the chemical specificity of protein-DNA interactions at residue and base-pair resolution, we studied the stability and folding pathways of a tetra-nucleosome. Dynamical simulations performed with an advanced sampling technique uncovered multiple pathways that connect open chromatin configurations with the zigzag crystal structure. Intermediate states along the simulated folding pathways resemble chromatin configurations reported from *in situ* experiments. We further determined a six-dimensional free energy surface as a function of the inter-nucleosome distances via a deep learning approach. The zigzag structure can indeed be seen as the global minimum of the surface. However, it is not favored by a significant amount relative to the partially unfolded, *in situ* configurations. Chemical perturbations such as histone H4 tail acetylation and thermal fluctuations can further tilt the energetic balance to stabilize intermediate states. Our study provides insight into the connection between various reported chromatin configurations and has implications on the *in situ* relevance of the 30nm fiber.

## Introduction

The organization for a string of nucleosomes is of fundamental interest and may dictate the transcriptional fate of enclosed genes.^1^ Significant research effort has been devoted to characterize chromatin structure *in vitro,*^2–4^ and has led to the discovery of the 30nm fiber.^5–8^ The precise arrangement of nucleosomes inside the fiber remains controversial, though high-resolution structures for an array of nucleosomes with well-defined positions are becoming available.^9,10^ Even less known is the dynamical process along which chromatin folds from extended configurations to these regular fibril structures.^11^ Methodologies that enable efficient conformational exploration and accurate stability evaluation are needed for a detailed characterization of chromatin structure and dynamics. They may also provide insight into the *in vivo* relevance of the 30 nm fiber, which has been under debate after numerous failed attempts at finding similar motifs inside the nucleus with a variety of techniques.^12–16^

Experimentally probing chromatin organization at atomistic resolution remains challenging due to difficulties in precise chromatin engineering and assembly. Computational modeling offers an alternative and promising approach, and has played crucial roles in uncovering the possible topology of nucleosomes in fibril structures.^17,18^ For example, the two angle model proposed by Woodcock and coworkers^7^ was instrumental for interpreting electron microscopy images of chromatin and for providing a consistent description of various chromatin fiber models. ^19–21^ Schlick and coworkers developed a mesoscale model that explicitly describes the nucleosome core particle and its charge distribution, the histone tails, and the linker DNA.^2,22,23^ Applications of these models have resulted in numerous insights into the sensitivity of chromatin organization with respect to nucleosome repeat length, linker histones, salt concentrations, etc.^24–28^ To better characterize the impact of DNA sequences, the Wilson group,^29,30^ Zhurkin group^31–34^ and van Noort group^35^ have developed chromatin models in which the DNA molecule is represented at a single base-pair resolution. The de Pablo group further introduced a systematic parameterization and coarse-graining procedure to derive mesoscopic models that are potentially of improved accuracy. ^36^

Here, we applied a near atomistic model to study the basic unit of chromatin fibers, the tetra-nucleosome. Protein and DNA molecules were represented with residue and base-pair resolution to provide a particle-based description of electrostatic interactions and specific protein-protein interactions that are crucial at stabilizing inter-nucleosome contacts. Dynamical simulations of this model uncovered two folding pathways that differ in the ordering of nucleosome contact formation. While the sequential pathway connects open chromatin configurations to the compact zigzag structure^10^ with tri-nucleosome structures, the concerted pathway goes through conformations that resemble a *β*-rhombus. These folding intermediates bare striking similarity to chromatin configurations observed *in situ*.^16,37^ We then computed the free energy surface as a function of the six inter-nucleosomal distances with a neural network approach and confirmed the statistical significance of these pathways. The free energy surface further suggests that while the stacked, zigzag configuration resides in the global minimum, the folding intermediates bear comparable stability. Significantly, these intermediate configurations can be further stabilized by configurational entropy, histone modifications, and variation in the secondary structure of histone tails, highlighting the sensitivity of chromatin organization to thermal and chemical perturbations. Our study suggests that chromatin configurations observed *in situ* might form as a result of local excitations or unfolding from the global minimum of the *in vitro* fibril structures.

## Results

### Direct simulations of tetra-nucleosome folding

We applied a near-atomistic model to characterize the folding process during which chromatin transitions from extended configurations with minimal contacts between nucleosomes to the compact zigzag crystal structure (PDB ID: 1ZBB).^10^ The near-atomistic model has been used extensively in prior studies to investigate nucleosome unwinding and nucleosome-nucleosome interactions (see *Methods*). A tetra-nucleosome with 20-bp-long linker DNA representing a minimal stable structural unit of chromatin fibers^9,11^ was employed in our simulations.

To facilitate observations of large-scale conformational rearrangements, we combined meta-dynamics^38^ with temperature-accelerated molecular dynamics (TAMD)^39,40^ to bias simulation trajectories along two collective variables, *Q* and *R_g_*. The fraction of native contacts *Q* is defined as

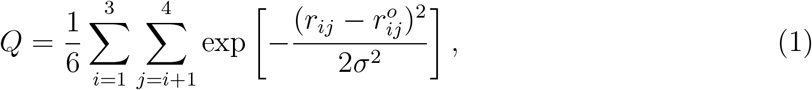

where *i* and *j* are indices of nucleosomes and *σ* = 2 nm. *r_i_j* measures the distance between the center of nucleosome *i* and *j*, and 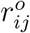 is the corresponding value in the crystal structure.^10^ *Q* measures the similarity between a given tetra-nucleosome configuration and the crystal structure, while *R_g_*, the radius of gyration, quantifies the overall compaction and size of the structure. As shown in an example trajectory (Fig. 1A), the tetra-nucleosome undergoes multiple collapse/expansion transitions, as indicated by Q varying from 0.1 to 0.8, and *R_g_* from 15 to 5 nm. On the other hand, without the advanced sampling techniques, tetra-nuclesome unfolding occurs on a much slower timescale and was not observed during the simulation time (Fig. S1).

**Figure 1:**
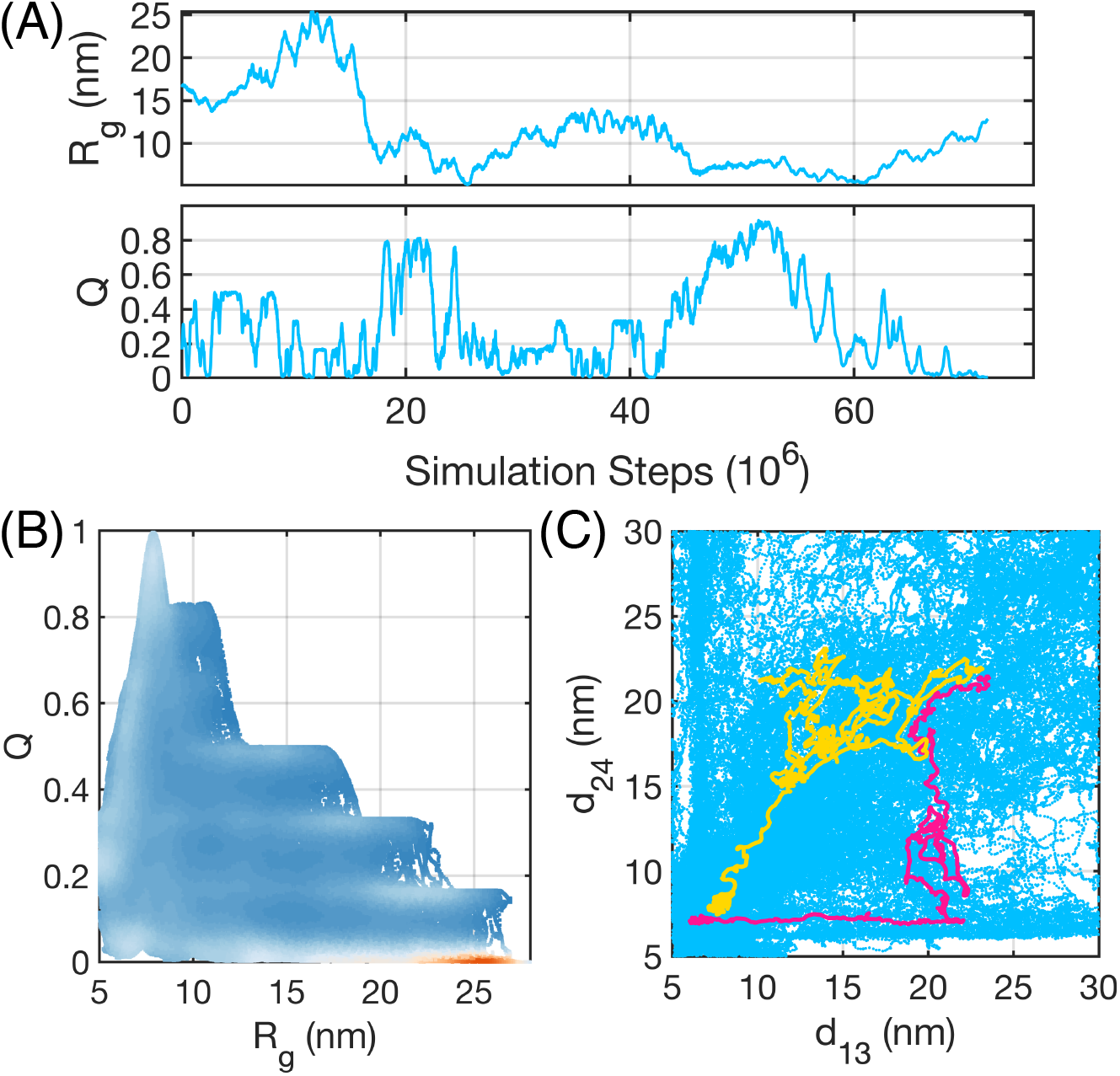
Direct simulations of the tetra-nucleosome folding process enabled via an advanced sampling technique. (A) Time evolution of the two collective variables along an example trajectory during which chromatin undergoes multiple folding and unfolding transitions. (B) Scatter plot of all the sampled tetra-nucleosome configurations represented by *R_g_* and *Q*. Color scale represents frequency estimated from direct counting that increases from blue to white and to red. (C) Scatter plot of all the tetra-nucleosome configurations represented by 1-3 and 2-4 nucleosome distances. Two example trajectories that correspond to the concerted and sequential folding pathway are highlighted in yellow and pink, respectively.

We carried out a total of 20 independent simulations starting from different open tetranucleosome conformations for a comprehensive exploration of the configurational space. Time evolutions of *Q* and *Rg* along these trajectories are provided in Fig. S2. Similar to the example shown in Fig. 1A, large scale changes in tetra-nuclesome conformations can be observed. Collectively, these trajectories provide a rather complete and relatively uniform coverage of the phase space (see Fig. 1B). We note that the negative correlation between *R_g_* and *Q* restricts the data points to the lower bottom of the plane. We further projected the simulated trajectories onto the distances between 1-3 and 2-4 nucleosomes, both of which reach their minimal value in the zigzag configuration. As shown in Fig. 1C, these two degrees of freedom are sufficiently sampled as well. The relatively complete coverage over these two new collective variables supports the efficacy of the employed sampling techniques for conformational exploration.

To gain more intuition into the simulated chromatin configurations, we used the K-means algorithm to identify the 100 most populated clusters. Example cluster central structures are shown in Fig. 2, and the full list is provided in Movie S1. Stacked configurations, for which the 1-3 and 2-4 nucleosomes are in close contact as in the crystal structure, can be readily seen. However, compared to the crystal structure, these conformations appear more compact and less twisted, with a decrease in the angle between vectors formed by 1-3 and 2-4 nucleosomes. Consistent with prior studies, histone H3 (orange) and H4 (purple) terminal tails were found to mediate the close contact between nucleosomes. ^23,41^ While H4 tails primarily bridge non-neighboring nucleosomes, H3 tails form contacts with linker DNA and bring neighboring nucleosomes close.

**Figure 2:**
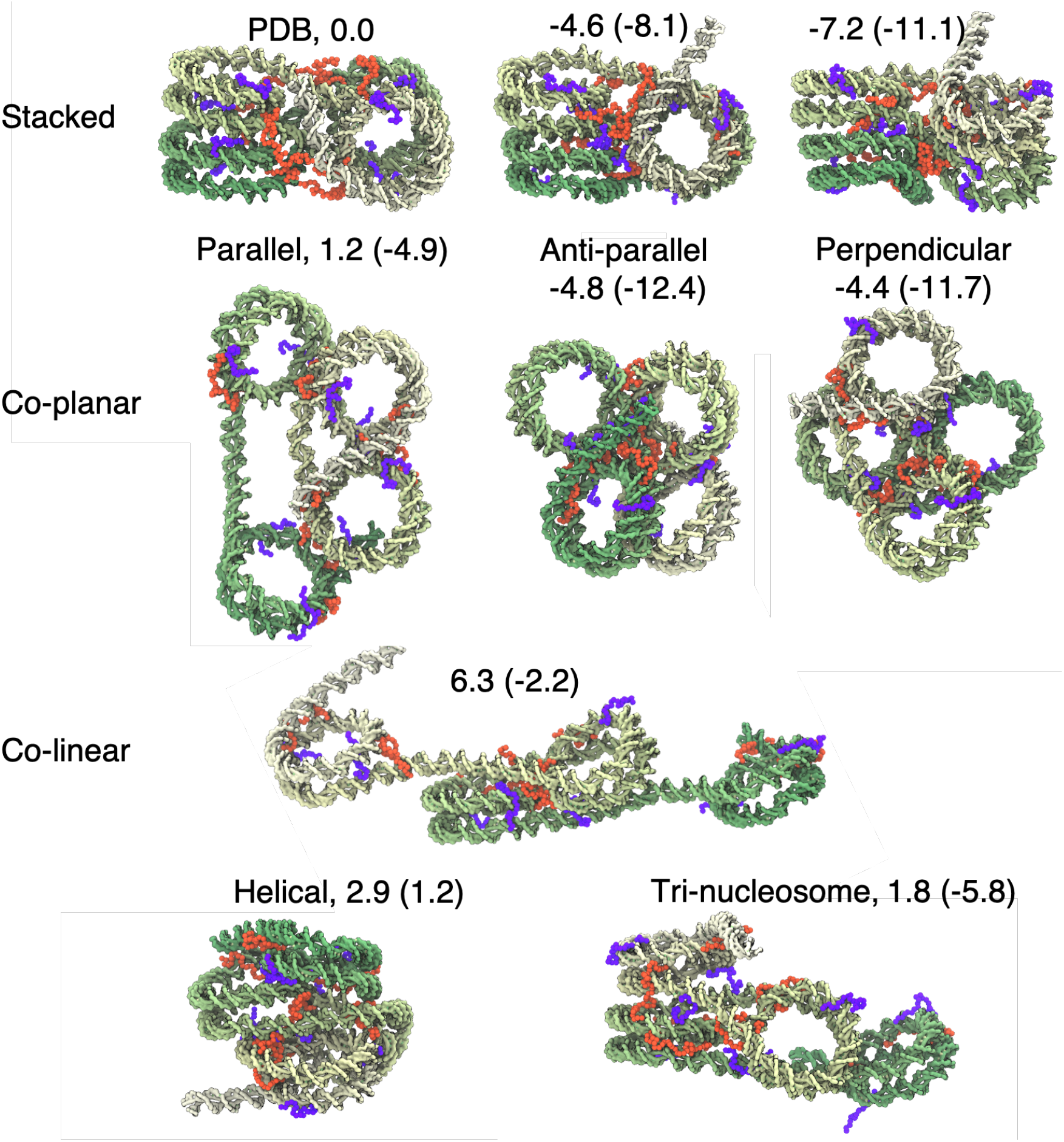
Representative tetra-nucleosome configurations from K-means clustering. See Movie S1 for the full set of cluster structures. For clarity, only the DNA molecule, the color of which varies from green to white, is shown. The N-terminal tails of histone H3 and H4 protein are drawn in purple and orange, respectively. The numbers indicate their stability relative to the PDB structure (top left) measured in the unit of kcal/mol, and were evaluated using simulations performed with the presence of protein-protein interactions between nucleosomes. The corresponding values inside the parentheses were obtained using simulations performed without such interactions.

In addition to the stacked configurations, we observed co-planar structures in which the two vectors formed by neighboring nucleosomes are either parallel, anti-parallel, or perpendicular to each other. There are also structures in which the four nucleosomes align almost along a straight line (co-linear) and along a left-handed helical path (helical). Finally, for trinucleosome structures, three nucleosomes adopt positions seen in the crystal structure, while the last one extends out. The disordered histone tails were again found to localize at the interface between nucleosomes to mediate contacts. Many of these structures are strikingly similar to those reported in prior studies that probe chromatin configurations *in situ*.^15,16^ Therefore, we anticipate that these structures could be useful for interpreting experimental results and provide their 3D coordinates on the group GitHub page (https://github.com/ZhangGroup-MITChemistry/TetraNucl).

### Constructing high-dimensional free energy surface with deep learning

The diversity of structures suggests that while *R_g_* and *Q* are sufficient for conformational exploration, additional collective variables are needed to fully capture the complexity of tetra-nucleosome organization. Furthermore, it would be beneficial to quantify the thermodynamic stability of the intermediate configurations. Due to the biases introduced in meta-dynamics and TAMD, such information cannot be directly obtained from the dynamical simulations.

We, therefore, determined the free energy surface as a function of the six inter-nucleosome distances, **d** = (*d*_12_, *d*_13_, *d*_14_, *d*_23_, *d*_24_, *d*_34_). These distances are crucial for differentiating various tetra-nucleosome configurations. We note that a high-dimensional free energy surface is challenging to calculate with traditional techniques, including umbrella sampling,^42^ due to their exponential increase in computational cost with dimensionality. Instead, we adopted a neural network approach to calculate the free energy from mean forces determined at a total of 10000 preselected centers^43^ (see Fig. 3A and *Methods* for details). A key advantage of this approach is that no overlap among centers is required, thus alleviating the curse of dimensionality by avoiding a uniform coverage of the phase space.

**Figure 3:**
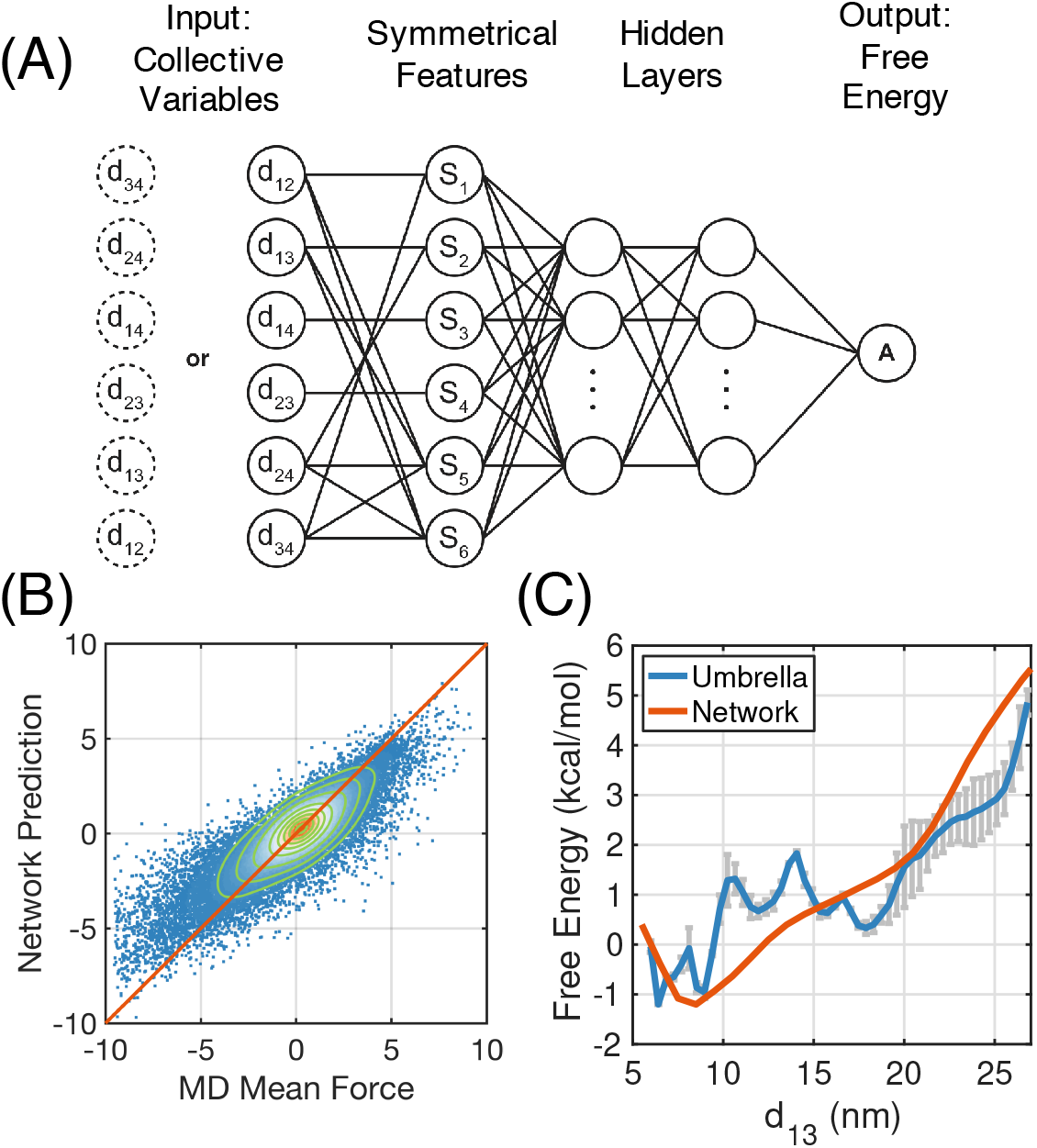
Parameterization and validation the six-dimensional free energy surface with a multilayer neural network. (A) Illustration of the neural network designed to take into account the symmetry of the structure with respect to index ordering. Input of this network is tetra-nucleosome configurations represented via inter-nucleosomal distances and output is the corresponding free energy. (B) Comparison between mean forces in the unit of kcal/mol/nm determined from restrained molecular dynamics (MD) simulations and neural network prediction. The density of data points increases as the color changes from blue to white to red. The green lines are constant density contours and the red diagonal line is shown as a guide for the eye. (C) Comparison between the free energy profile as a function of 1-3 nucleosome distance determined from neural network (orange) and umbrella sampling (blue). The errorbars represent one standard deviation from the mean.

Once parameterized, the neural network can provide free energy estimations for any configuration represented by the six inter-nucleosome distances. Mean forces can be determined from the network as well via the back-propagation algorithm. ^44^ As shown in Fig. 3B, the mean forces calculated using the trained neural network are in good agreement with the ones produced from restrained molecular dynamics simulations, and the Pearson correlation coefficient between the two is 0.87. While fine-tuning hyperparameters can lead to higher correlation coefficients, the network becomes less robust and tends to overfit the data (see Figs. S3 and S4), partially due to the inherent uncertainty in mean force estimations using molecular dynamics (MD) simulations.

To directly validate the accuracy of the free energy estimation, we integrated over the other five degrees of freedom to obtain a one-dimensional free energy profile as a function of the distance between 1-3 nucleosomes. For comparison, we carried out additional simulations to compute the same quantity using umbrella sampling^42^ and the weighted histogram analysis method (WHAM).^45^ Details for these simulations are provided in the supporting information (SI). As shown in Fig. 3C, despite the significant difference between the two methodologies, the two free energy profiles agree well with each other, and deviations between them are less than 1 kcal/mol across the entire range of distance. We note that the free energy profile from umbrella sampling is rather rugged, despite our significant simulation effort (Table S1). The ruggedness arises because the system at neighboring umbrella windows can be trapped in different tetra-nucleosome configurations with comparable *d*_13_ and interconversion among these configurations is slow.

We next used the neural network to quantify the stability of the example structures shown in Fig. 2. The free energy values are provided along with the structures in parentheses. We found that stacked conformations with more aligned nucleosome columns have lower free energy than the PDB structure. For example, the 1-3 nucleosome axis is more parallel to the 2-4 nucleosome axis in these conformations. Rotating the two axes relative to each other could cause twisting in the linker DNA, potentially explaining the higher free energy of the PDB structure. The rotation in the PDB structure could be a result of crystal packing as well as the high concentration of Mg^2+^ ions used in sample preparation.^10,46,47^ To ensure that the predicted structural change is not an artifact of the near-atomistic model used here, we carried out additional simulations using the SIRAH force field.^48^ While molecules are still treated at a coarse-grained level, this force field includes explicit representations of water molecules and ions and is expected to describe the solvation effect and electrostatic interactions accurately. As shown in Fig. S5, both an unbiased 4.0-μs-long dynamical trajectory and the free energy profile estimated from a series of ~1.0-μs-long umbrella sampling simulations support the stability of the less rotated conformations.

In addition, we found that some of the co-planar structures adopt comparable, or even lower, free energy values relative to the stacked configurations. Their stability potentially arises from the absence of close contacts between inter-nucleosomal DNA seen in the stacked structures while maintaining favorable interactions between histone tails and the DNA. One or both of these two features are missing or compromised in co-linear, helical, and trinucleosome configurations, giving rise to their higher free energy. We note that the computed free energy is a function of inter-nucleosome distances and measures the stability for an ensemble of structures, not just the ones shown in Fig. 2. For example, individual nucleosomes and histone tails can rotate and diffuse without significantly altering inter-nucleosome distances. These movements further contribute to configurational entropy and the stability of the co-planar structures as well but are severely impaired in the stacked configurations due to their tight packing.

### Identifying most probable chromatin folding pathways

The high dimensional free energy surface also enables a rigorous characterization of the folding pathways to form stacked structures. Such pathways could offer insight into the stability of regular fibril configurations and their relationship to the more irregular structures often seen *in situ*. Our exploratory, dynamical simulations indicated a complex mechanism with multiple folding pathways, as highlighted by the two continuous trajectories in Fig. 1 (see Movies S2 and S3). The formation of contacts along the yellow trajectory is more concerted, and there is a strong correlation between the two non-neighboring inter-nucleosome distances. On the other hand, 1-3 nucleosomes remain far apart as 2-4 nucleosomes stack on top of each other along the pink, sequential trajectory. However, since these trajectories were obtained from biased simulations, their thermodynamic relevance remains to be shown.

Starting from the parameterized free energy surface, we searched for the most probable chromatin folding pathways that connect an extended configuration with the stacked conformation (Fig. 4A) using the finite-temperature string method^49,50^ (see *Methods* Section for details). As shown in Fig. 4B and Fig. S6, three sets of distinct pathways were identified. Consistent with the exploratory trajectories shown in Fig. 1C, two of the paths indeed correspond to the sequential pathway, and the third one represents the concerted pathway. Also, tetra-nucleosome configurations sampled along the paths are well separated, supporting the statistical significance of the difference between them. For simplicity, the paths were shown in the two dimensions of 1-3 and 2-4 inter-nucleosome distances. The variations of all distances along the paths are provided in Fig. S7. Our observation of sequential and concerted pathways in both string method calculations and dynamical simulations supports their robustness and the independence of the folding mechanism from our choice of collective variables and computational techniques.

**Figure 4:**
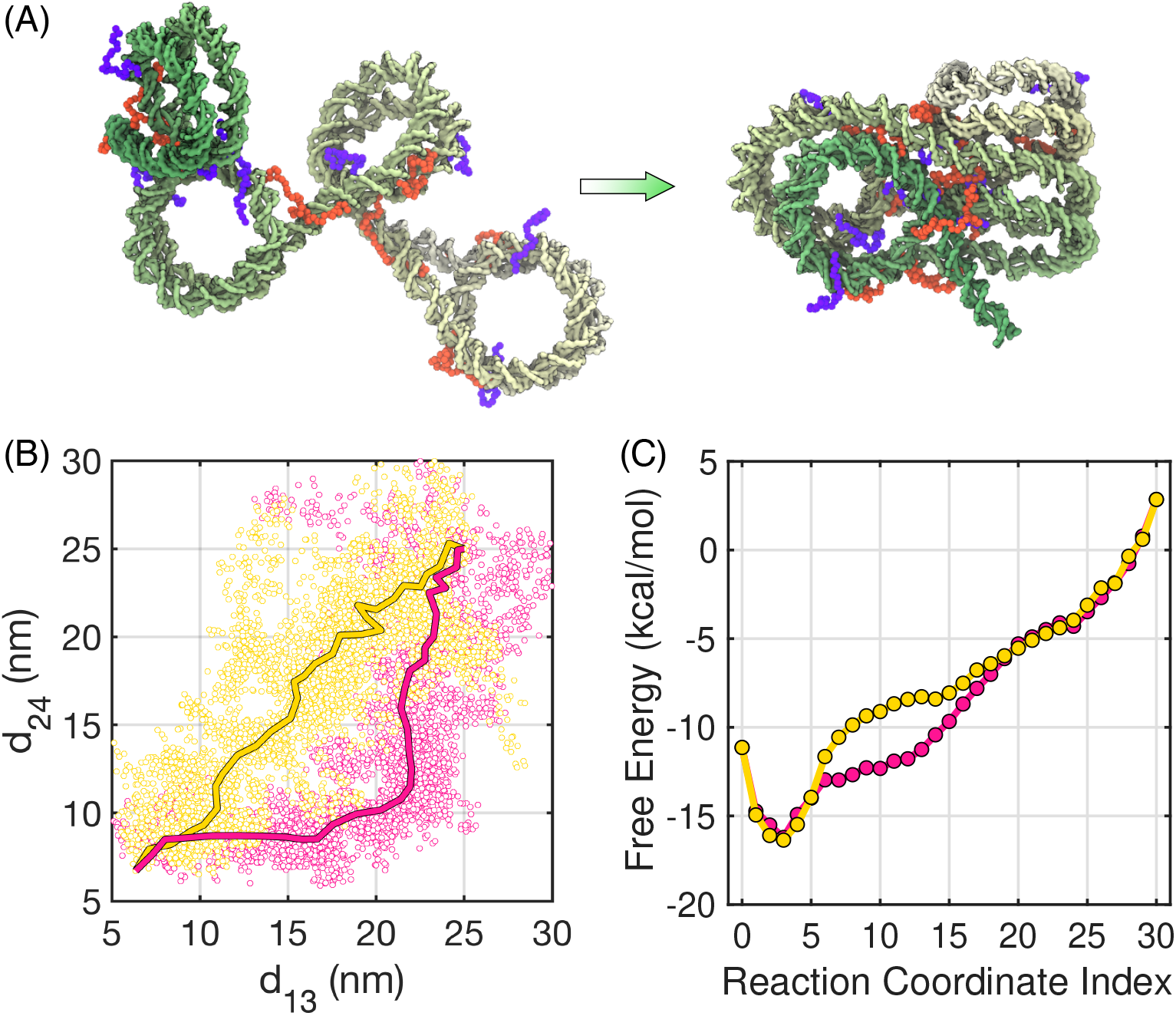
Chromatin folding pathways determined using the finite temperature string method. (A) Illustration of the starting and end configurations of the chromatin folding pathways using the same coloring scheme as in Figure 2. (B) The sequential (pink) and concerted (yellow) pathways plotted over the two inter-nucleosome distances. The dots correspond to tetra-nucleosome configurations obtained from individual simulations restricted to Voronoi cells defined using images along the path. They represent fluctuations around the path on the scale of *k_B_T*. (C) Free energy profiles along the two pathways. See Fig. S6 for corresponding results of the second sequential pathway.

Next, we reconstructed chromatin configurations along the calculated strings using restrained molecular dynamics simulations centered at the corresponding inter-nucleosome distances. Upon close examination, we found that the example structures shown in Fig. 2 emerge as intermediates along the folding pathways. In particular, the concerted path goes through co-planar configurations that resemble parallel or perpendicular structures (see Movie S4). Similarly, tri-nucleosome structures can be readily seen along the sequential pathway (Movie S5). Therefore, these chromatin structures observed by *in situ* experiments can be viewed as *en route* to fibril conformations. We did not observe anti-parallel structures despite their apparent stability. Further investigations suggest that it might serve as a kinetic trap for forming fibril structures and must be undone before folding proceeds (see Fig. S8).

In Fig. 4C, we plotted the free energy profiles along the representative sequential and concerted pathways shown in Movies S4 and S5. These profiles compare favorably to the force-extension curves obtained by van Noort and coworkers using single-molecule force spectroscopy experiments^35,51^ (see Fig. S9). An almost continuous decrease of the free energy supports chromatin folding as downhill transitions. The similarity of the two profiles argues for the significance of both pathways with comparable kinetics. The sequential pathway does appear to transition across more favorable chromatin configurations with lower free energy starting at reaction coordinate index 16. Such favorability arises because contacts between nucleosomes are formed earlier in the tri-nucleosome structures than in the intermediates along the concerted string.

### Determinants of chromatin conformational stability

The quantitative free energy surface suggests that the fibril configurations with stacked nucleosomes are not particularly stable compared to partially open, unfolded structures. Their weak stability can also be seen from the relative flatness of the free energy profile as a function of 1-3 and 2-4 nucleosome distances around the global minima (Fig. 5A). In the following, we investigate in detail factors that contribute to the stability of chromatin organization.

**Figure 5:**
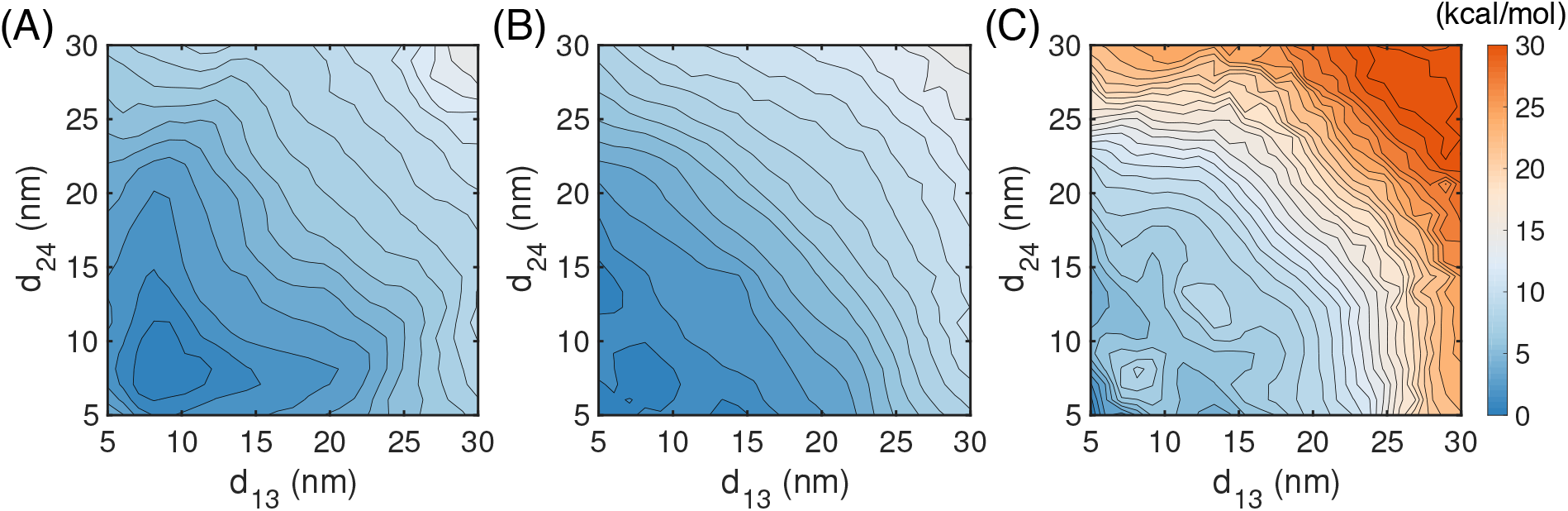
Impact of specific and non-specific interactions on chromatin stability. (A,B,C) Contour of two dimensional free energy profile as a function of the distance between 1-3 (*d*_13_) and 2-4 (*d*_24_) nucleosomes at 1 kcal/mol interval for the system without (A) and with (B) protein-protein interactions at 300 K and with protein-protein interactions at 250 K (C).

Histone proteins from different nucleosomes may form contacts to promote chromatin folding and stabilize the zigzag crystal structure. Therefore, we introduced specific interactions between proteins using the Miyazwa-Jernigan (MJ) potential that has been widely used for studying protein folding. ^52,53^ A new set of simulations were carried out to estimate mean forces with the presence of protein-protein interactions, starting from the same set of configurations as those used for results shown in Fig. 3. We then trained another neural network to compute the free energy surface from these mean forces. As shown in Figs. 5B, the global minimum indeed shifts toward smaller inter-nucleosome distances. Notably, chromatin configurations with stacked nucleosomes are now favored over the co-planar structures (see Fig. 2). To our surprise, the overall quantitative change on the free energy profile from the one computed without accounting for protein-protein interactions appears minimal, suggesting the importance of additional factors for chromatin stability.

We next studied, in addition to specific interactions, the impact of non-specific contributions from configurational entropy on chromatin stability. We repeated the mean force calculations with protein-protein interactions, but at a temperature of 250 K instead of 300 K. As shown in Figs. 5C, the free energy surface parameterized from these mean forces strongly favors the crystal structure as the most stable configuration. This result suggests that the stability of unfolded, irregular chromatin structures, at least in part, arises from the configurational entropy. As mentioned before, unlike the stacked configurations, these structures place less constraint on the movement of individual nucleosomes and histone tails and are expected to be more dynamic.

Finally, since histone proteins undergo a variety of post-translational modifications that affect inter-nucleosome interactions, we explored the impact of acetylation of the H4 tail on chromatin stability. We first focused on the residue K16, the acetylation of which has implications in aging. ^54^ Specifically, we neutralized the positive charge on the residue to mimic the chemical effect of acetylation, repeated the mean force calculations without specific protein-protein interactions at 300K, and retrained the neural network to compute the free energy. As shown in Fig. 6A, the resulting free energy profile is somewhat similar to the wild type result shown in Fig. 5A with a different color scale. Prior studies have suggested, in addition to electrostatic interactions, acetylation could alter the secondary structure of histone tails.^23,55,56^ Via a multi-scale approach, Collepardo-Guevara et al. further showed that an increase in structural ordering of H4 tail could unfold chromatin by limiting internucleosomal interactions. ^23^ Therefore, we carried out additional simulations that both remove the positive charge on K16 and restrain the H4 tail to a beta-sheet motif uncovered in all-atom explicit solvent simulations. ^23^ As shown in Fig. 6B, consistent with the results by Collepardo-Guevara et al., unfolded chromatin configurations become more stable, and the free energy profile exhibits additional basins at large inter-chromosomes distances. Therefore, for the single residue acetylation, the impact of secondary structure changes appears to outweigh electrostatic interactions. We further studied an additional system in which the entire H4 tail is acetylated to evaluate the overall contribution of all the nine positive charges to chromatin stability. To focus on the role of electrostatic interactions, we did not introduce biases to the tail secondary structure. The resulting two-dimensional profile (Fig. 6C) again supports the stability of open chromatin conformations with large inter-nucleosome distances. Therefore, charged interactions, especially when cumulated over multiple residues, contribute to H4 tails’ role in stabilizing compact chromatin conformations as well.^24,57^

**Figure 6:**
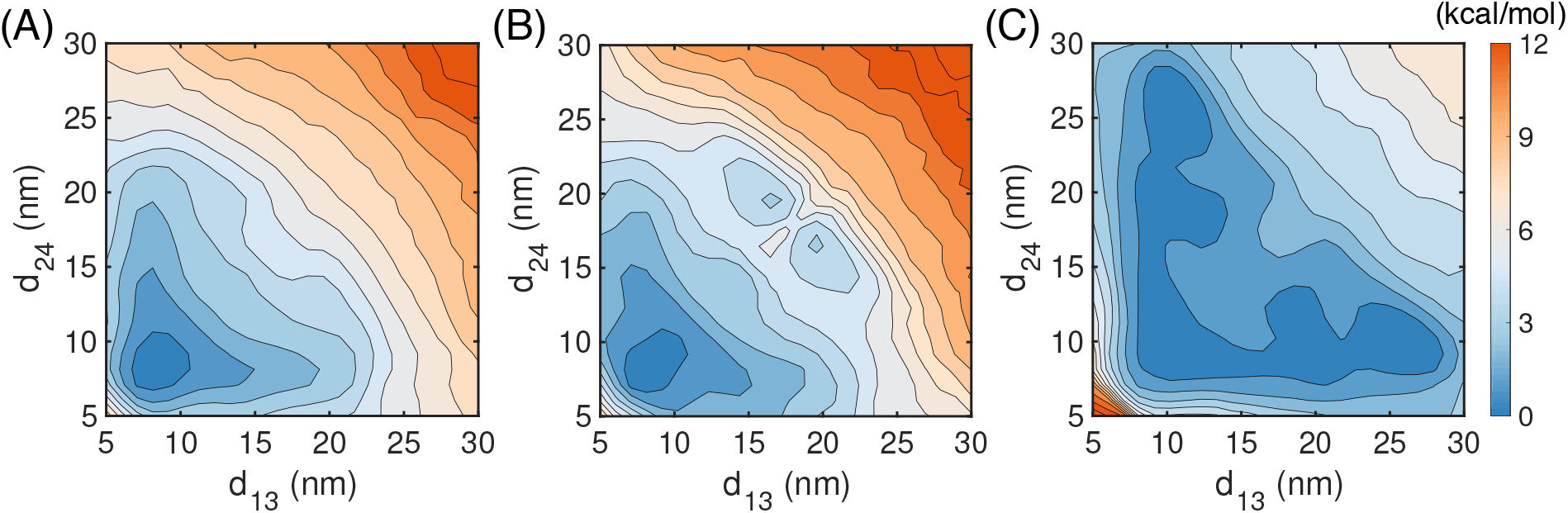
Impact of histone H4 tail acetylation on the free energy landscape of chromatin folding. (A,B,C) Contour of two dimensional free energy profile as a function of the distance between 1-3 (*d*_13_) and 2-4 (*d*_24_) nucleosomes at 1 kcal/mol interval for the system that removes the positive charge on K16 (A), that removes the positive charge on K16 and biases the tail to a beta-sheet conformation (B), and that removes all the nine positive charges on the tail (C).

## Conclusions and Discussion

We studied tetra-nucleosome folding with a near-atomistic force field that models protein and DNA molecules at single residue and base-pair resolution. Compared to mesoscopic models, this force field gives rise to a more rugged energy landscape and a significant increase in computational cost, creating challenges for chromatin conformational sampling. We found that a technique that combines meta-dynamics with TAMD succeeded in producing continuous trajectories that connect open, extended structures to collapsed and folded ones. A deep-learning approach, together with mean force calculations, further allowed the computation of a six-dimensional free energy surface around the phase space explored by the biased simulations. The free energy surface provides unbiased evaluation on the relative stability of tetra-nucleosome configurations and its sensitivity to chemical and thermal perturbations.

Our simulations revealed a wide variety of 3D configurations that tetra-nucleosomes could adopt in addition to the zigzag structure resolved by X-ray crystallography. ^10^ Several of these simulated configurations have indeed been reported in prior experimental studies. In particular, by performing cryo-electron tomography on a HeLa cell, ^15^ Gan and coworkers directly observed tri-nucleosome configurations similar to the one shown in the bottom of Fig. 2, and found that they are more prevalent than compact structures consisting of four or more nucleosomes. On the other hand, structural motifs that resemble the parallel (*β*-rhombus) and helical (*α*-tetrahedron) have been reported in models derived from chromosome conformation capture experiments that measure spatial proximity between nucleosomes inside the nucleus via chemical cross-linking and high-throughput sequencing. ^16^

The relationship between *in situ* chromatin configurations and the zigzag crystal structure is evident from the dynamical trajectories shown in Fig. 1C (yellow and pink) and Movies S2 and S3. These trajectories continuously bridge tri-nucleosome and *β*-rhombus configurations with the zigzag structure as time progresses. Therefore, the *in situ* configurations can be viewed as folding intermediates along the pathways towards the zigzag structure. We note that the dynamical trajectories were obtained from simulations biased over two collective variables, *R_g_* and *Q*. Mechanistic insights drawn from these simulations are only reliable if the two variables represent the slowest degrees of freedom of the system. Otherwise, they may obscure the importance of true reaction coordinates that dictate chromatin folding. Importantly, however, string method calculations that track the most probable pathways for tetra-nucleosome folding support the same conclusions as the dynamical simulations. These calculations used a larger set of collective variables and an unbiased free energy surface. They provide strong support for the thermodynamic significance of the folding mechanism observed in exploratory trajectories.

Our study has significant implications on the *in situ* relevance of the 30 nm fiber. ^58,59^ The free energy surfaces support the global stability of regular, fibril configurations for isolated chromatin. Since *in situ* configurations correspond to folding intermediates, they can also be viewed as local excitations or unfolding from the global minimum. Such excitations are not too rare as the energetic penalty arising from a loss of inter-nucleosome contacts can be largely compensated by an increase in the configurational entropy. Histone modifications, and potentially other factors such as linker length, can even tilt the balance to drive the global stability towards intermediate configurations. Given the heterogeneous environment inside the nucleus, it is perhaps not too surprising to find the prevalence of partially unfolded, irregular structures.

As aforementioned, the near-atomistic model remains computationally costly for simulating long chromatin segments. Therefore, we anticipate that it will be most effective for simulating systems of tens of nucleosomes in size and evaluating the relative stability of various topological forms.^17,18,60,61^ For studying conformational arrangement of genomic regions that are 10~100 kb in length, mesoscopic models introduced by the Schlick or the de Pablo group offer a more promising route with better balanced accuracy and computational efficiency. ^22,62^ At even larger scales such as a whole chromosome or the entire genome, integrative approaches that take into account the impact of the nucleus environment with experimental constraints may provide more faithful structural models for chromatin. ^63,64^

Several aspects of the near-atomistic model can be further improved for better accuracy. In particular, fine-tuning protein-protein interactions could lead to a more balanced treatment of both ordered and disordered regions of histone proteins. Introducing base pair specific protein-DNA interactions may help account for DNA sequence-dependent chromatin stability.^65^ Furthermore, our adoption of the Debye-Hückel theory and implicit ions may be insufficient for electrostatic interactions, especially when considering the impact of divalent ions such as Magnesium.^66,66^ Finally, as shown here and by Collepardo-Guevara et al.,^23^ histone tail secondary structure changes upon post-translational modifications can impact chromatin stability as well and must be approached with caution.

## Methods

### Near-atomistic modeling of chromatin organization

We applied a near-atomistic model to study chromatin with increased chemical details. One bead per amino acid and three sites per nucleotide were employed to describe protein and DNA molecules, respectively, leading to a system of 8058 coarse-grained beads in size. Following the same procedure as in structure-based models, ^67,68^ contact potentials derived from the crystal structure^10^ were introduced for amino acids within individual histone octamers to stabilize the tertiary structure. No specific interactions between proteins from different nucleosomes were included unless otherwise specified. The 3SPN2.C force field developed by de Pablo and coworkers ^69–71^ was used to model the DNA molecule. Protein-DNA interactions were described with the Debye Hückel potential at a salt concentration of 150 mM and the Lennard-Jones potential for excluded volume effect. More details on the model setup and force field parameters can be found in the SI.

We anticipate the improved resolution compared to mesoscopic models to be beneficial at modeling electrostatic and protein-protein interactions within and among nucleosomes. Similar models have been used extensively to study the dynamics and stability of single nucleosomes.^41,72–76^ They were shown to reproduce the energetic cost of nucleosomal DNA unwinding, ^72,74^ the dependence of the energetic cost on applied tension, ^73^ and the sequencespecific DNA binding strength to the histone octamer.^77^ The de Pablo group recently showed that near-atomistic modeling could reproduce the binding strength between a pair of nucleosomes measured in DNA origami-based force spectrometer experiments. ^41,78^

To further evaluate the force field’s accuracy on connected nucleosomes, we simulated a pair of di-nucleosomes connected with linker DNAs of 50 and 55 bp in length and monitored the unwrapping of nucleosomal DNA. As shown in Fig. S10, these simulations succeeded in capturing the subtle difference between the two systems. In particular, consistent with Förster resonance energy transfer (FRET) experiments,^79^ simulations predicted a more significant unwrapping for the di-nucleosome with 55-bp-long linker DNA. The populations of the unwrapped state for the two di-nucleosomes are in good agreement with experimental values as well.

### Simulation details

To improve the near-atomistic model’s efficiency for chromatin simulations, we grouped the ordered part of histone proteins and the inner 73 bp of nucleosomal DNA together as rigid bodies (see Fig. S11A). Positions and velocities of all the atoms within each rigid body were updated together such that the body moves and rotates as a single entity. Disordered histone tails, outer nucleosomal DNA, and linker DNA remained flexible and no restrictions were applied on their conformational dynamics. Our partition of the rigid and flexible parts is motivated by findings from prior studies. In particular, the outer DNA does not bind tightly to histone proteins and can unwind spontaneously, ^80^ while the inner layer is much more stable, and its unwinding is prohibited by a sizeable energetic barrier. ^74,81^ Furthermore, the histone core remains relatively stable during DNA unwinding.^72,82^ As shown in Fig. S11B, the rigid body treatment has minimal impact on the model’s accuracy in simulating internucleosome distances.

Molecular dynamics simulations were carried out with the LAMMPS software package. ^83^ A time step of 5 fs was used for configurational exploration and 10 fs for mean force calculations. We note that the near-atomistic model’s timescale is not carefully calibrated, and one step here can correspond to a much longer time step in all-atom simulations. ^84^ The Nose-Hoover thermostat was applied separately to the rigid and flexible parts of the system to maintain the simulations at a temperature of 300 K with a damping coefficient of 1 ps. All simulations were carried out in a cubic box with sides of 200 nm.

### Driving chromatin folding with advanced sampling techniques

To explore the folding landscape of the tetra-nucleosome and collect a diverse set of conformations, we combined metadynamics with TAMD to bias the simulations along two collective variables *R_g_* and *Q* using PLUMED.^85^ While meta-dynamics enhances barrier crossing by introducing memory dependent potentials to penalize the system from revisiting the sampled region of phase space, TAMD accelerates rare transitions with fictitious dynamics of collective variables performed at high temperatures. Combining the two techniques has been shown to result in superior efficiency for conformational sampling compared with simulations conducted using only one of them. ^86^ Implementation details of the biases can be found in the SI.

We carried out 20 independent biased simulations. A total of at least 50 million steps were carried out for each simulation (see Fig. S2), and we saved the configurations along the trajectories every 5000 steps. In total, 293291 tetra-nucleosome configurations were recorded. Initial configurations of these simulations were sampled from a 40×10^6^ step long trajectory in which the tetra-nucleosome was restricted to open configurations with 1-3 and 2-4 nucleosome distances larger than 30 nm.

### Learning the six dimensional free energy surface using neural networks

We identified 10,000 representative tetra-nucleosome configurations from the biased exploratory trajectories using the *K*-means clustering algorithm. We then carried out additional restrained molecular dynamics simulations to estimate the mean forces ^87^ at each one of these representative configuration, **d**_o_, as

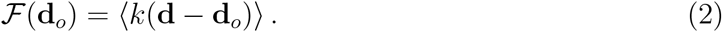

The angular bracket represents averaging over the structural ensemble derived from the energy function of the restrained tetra-nucleosome 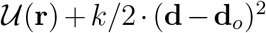. Finally, we parameterized the free energy surface 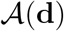 using a neural network that minimizes the difference between its gradient 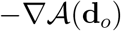 and the mean forces 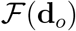. We further took advantage of the underlying symmetry of the system when designing the neural network. In particular, since the free energy is a function of internal coordinates, it should be independent of the ordering and indices of the nucleosomes. Therefore, the following two sets of distances, **d** = (*d*_12_, *d*_13_, *d*_14_, *d*_23_, *d*_24_, *d*_34_) and 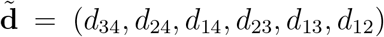, obtained from reordering the indices of the nucleosomes, should have approximately the same free energy, i.e., 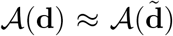. They are approximately equal because the DNA sequence used in our simulations is not palindromic. To enforce the symmetry in the neural network’s output, we converted the distances into symmetrical features such that 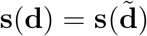 and parameterized the free energy as a function of **s** (Fig. 3A). More details on the architecture and parameterization of the neural network can be found in the SI.

### Uncovering folding pathways with the finite temperature string method

The finite temperature string (FTS) method ^50^ was used to determine chromatin folding pathways. The output of this method is a path (principal curve) that connects the reactant with the product while taking into account global features of the free energy surface and entropic effects of the transition. It is particularly useful when the free energy landscape is rugged, as is the case here, because it can average over fluctuations on the scale of *k_B_T* or smaller to reveal large scale characteristics of the landscape. The ruggedness is potentially an intrinsic feature of the tetra-nucleosome system, but it may also arise from statistical noise in mean force calculations and neural network parameterization. To avoid biases that might arise from initialization, we carried out a total of 70 string calculations starting from different initial path configurations for a comprehensive exploration of the free energy surface. Details on the implementation of the string method can be found in the SI.

## Data Availability

The datasets generated during and/or analysed during the current study are available in the GitHub repository (https://github.com/ZhangGroup-MITChemistry/TetraNucl).

## Code availability

The computer code used in this study is available upon reasonable request to the corresponding author.

## Acknowledgements

This work was supported by the National Institutes of Health (Grant 1R35GM133580). We thank Dr. Collepardo-Guevara for sharing the atomistic structure of histone H4 tail, Dr. van Noort for providing the FRET data on di-nucleosomes, and Dr. Pantano for sharing the simulation trajectory of tetra-nucleosome.

